# Functional partitioning of lipoic acid decouples cellular abundance from mitochondrial utilization

**DOI:** 10.64898/2026.05.22.727209

**Authors:** Pieter R. Norden, Riley J. Wedan, Abigail E. Ellis, Madeleine L. Hart, Megan R. Gendjar, Ryan D. Sheldon, Sara M. Nowinski

## Abstract

α-Lipoic acid (LA) is widely included in “mitochondrial cocktails” recommended to patients with primary mitochondrial disorders, yet its mechanism of action remains unclear. Here, we define the intracellular availability and functional utilization of LA in mammalian cells. We show that LA exists in two functionally distinct cellular pools: a low-abundance free pool and a protein-bound pool generated through mitochondrial fatty acid synthesis (mtFAS). Disruption of the mtFAS pathway abolishes protein lipoylation and impairs oxidative phosphorylation without altering free LA levels. Conversely, supplementation with exogenous LA markedly increases free intracellular LA without restoring protein lipoylation, mitochondrial respiration, or cell proliferation. Instead, the cellular effects of LA supplementation resemble those of the antioxidant N-acetylcysteine. These findings clarify the mechanism of action of a widely used mitochondrial supplement and identify a fundamental disconnect between cellular LA abundance and mitochondrial utilization, challenging the rationale for using LA supplementation to restore mitochondrial function.

## 1. Introduction

Mitochondrial dysfunction is a major therapeutic target in a wide range of diseases, including metabolic syndrome, inherited mitochondrial disorders, cardiovascular disease, neurodegeneration, and cancer^1,2^. A common pathological feature shared among these conditions is impaired mitochondrial oxidative phosphorylation (OXPHOS), which compromises cellular ATP production and increases reactive oxygen species (ROS) generation^1,2^.

Accordingly, pharmacological strategies aimed at restoring or boosting mitochondrial function and OXPHOS activity have become an important focus of therapeutic development^3–5^. Among these conditions, primary mitochondrial disease presents particular therapeutic challenges as patients present heterogeneously at diverse ages, display variable prognosis, and have increased risk of mortality at a young age^6^. The only therapies currently available are limited to non-curative symptom mitigation and often, despite treatment, conditions tend to worsen over time as patients age^7^.

Since no therapies to restore functional OXPHOS in primary mitochondrial disorders exists, patients are often recommended a “mitochondrial cocktail”, which consists of antioxidants and metabolic cofactors intended to bypass defects in the mitochondrial electron transport chain (ETC)^8,9^. A key component of this cocktail is α-lipoic Acid (often abbreviated aLA, or simply LA). LA is an essential cofactor that is covalently attached to several mitochondrial enzyme complexes which play key roles in central carbon metabolism. Most of these enzymes, collectively referred to as ‘lipoylated proteins’, are mitochondrial 2-ketoacid dehydrogenases, including pyruvate dehydrogenase (PDH), α-ketoglutarate dehydrogenase (OGDH), branched-chain ketoacid dehydrogenase (BCKDH), and 2-oxoadipate dehydrogenase (OADH), along with the H-protein of the glycine cleavage system (GCSH)^10,11^. The redox-active disulfide bond of LA provides the reducing equivalents required for catalysis by the dehydrogenases and contributes to their stabilization and redox-dependent regulation. Therefore, in the absence of lipoylation, these enzymes are catalytically dead ^10,12–14^.

As such an important mitochondrial co-factor, it is not surprising that LA is added to mitochondrial treatment cocktails in hopes of boosting mitochondrial function. However, eukaryotes lack a functional LA salvage mechanism, raising questions about how LA supplementation might be beneficial^10,15^. In fact, although several preclinical and clinical studies report improvements following LA supplementation, the underlying mechanisms remain poorly defined, and evidence for sustained clinical benefit is limited^16,17^. Beyond its role as an enzyme co-factor, free (not protein bound) LA has been proposed as a component of cell membranes^18^, and is known to act as a cell-permeant antioxidant^17,19,20^. However, the cellular source(s) of free LA and its abundance under normal culture conditions have not been rigorously defined. We therefore questioned whether and how free LA is produced in cells, how free LA levels correspond to protein-bound LA levels, and whether LA supplementation was able to confer cellular benefit(s) in the form of enhanced mitochondrial function, anti-oxidant effect(s), or other potential mechanisms.

Here, we show that cells have two distinct pools of LA that are functionally isolated: free LA, and LA covalently attached to lipoylated proteins. In cells unable to endogenously synthesize LA as a result of defects in the mtFAS pathway, protein lipoylation is near-completely absent, but free LA levels were not significantly altered across a variety of cellular models. Furthermore, we demonstrate that LA supplementation raises free intracellular LA, but can neither restore protein lipoylation, nor OXPHOS function, in mtFAS-deficient cells, revealing a fundamental disconnect between LA abundance and mitochondrial utilization. LA supplementation instead mirrored treatment with the common antioxidant N-acetylcysteine, perhaps suggesting that raising cellular levels of free LA may act predominantly via its antioxidant properties.

## 2. Results

### 2.1 Free cellular LA abundance is independent of mtFAS activity

To determine whether intracellular free LA abundance corresponded to levels of protein bound LA, we first assessed protein lipoylation status in mammalian cells unable to synthesize LA via the only known pathway, downstream of mitochondrial fatty acid synthesis (mtFAS). The mtFAS pathway builds fatty acyl chains that can reach up to 16 carbons in length on an acyl carrier protein (mtACP)^21^. LA biosynthesis branches off from the mtFAS pathway when an eight-carbon, fully saturated acyl-chain is built on mtACP (octanoyl-ACP)^21,22^. This octanoyl-intermediate is further modified to form LA and transferred directly to target proteins by a series of enzymes: lipoyl-transferase 2 (LIPT2), lipoic acid synthase (LIAS), and lipoyl-transferase 1 (LIPT1) ^10^.

Cells with mtFAS pathway deficiencies cannot synthesize octanoyl-ACP and therefore display protein lipoylation defects across a variety of species and cell types^21,23^. We verified this phenotype in murine skeletal myoblasts (C2C12), rat cardiomyoblasts (H9c2), and human epithelial adenocarcinoma (HeLa) cells. As predicted, we found that lipoylation of PDH and OGDH subunits (DLAT and DLST) was nearly absent in C2C12 deficient for the mtFAS enzymes malonyl-CoA-acyl carrier protein transacylase (MCAT) and mitochondrial trans-2-enoyl-coenzyme A reductase (MECR)(**Fig. 1A**). Similarly, MECR-deficient HeLa cells and MCAT-deficient H9c2 also exhibited marked reductions in lipoylation of PDH and OGDH subunits compared to clonal controls (**Fig. 1A**). Together, this confirms that mtFAS activity is required for LA biosynthesis and protein lipoylation across mammalian cell types.

**Figure 1.**
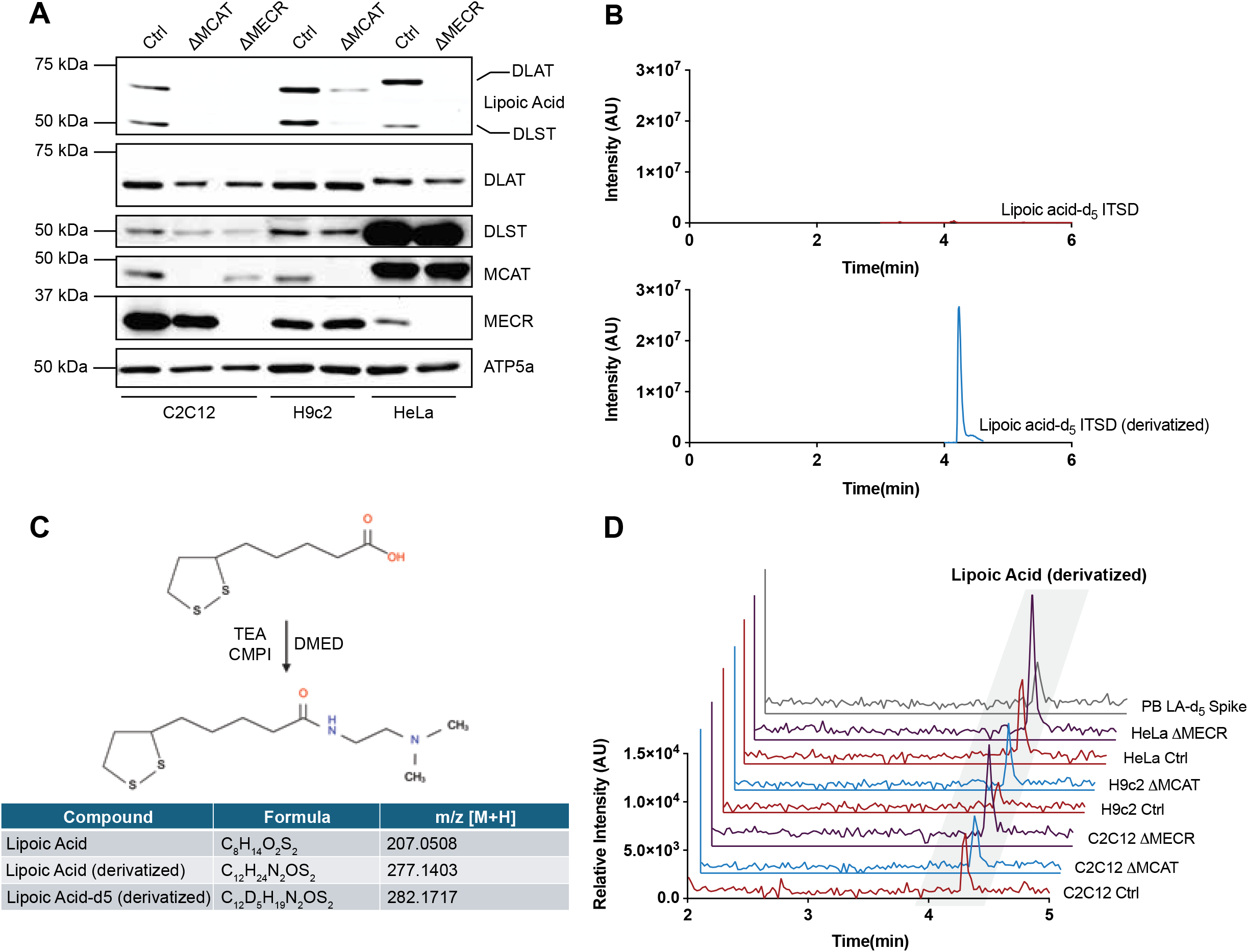
Free cellular LA abundance is independent of mtFAS activity. (A) Whole cell lysates or crude mitochondrial extracts were separated by SDS-PAGE and immunoblotted for the denoted targets in clonal control and MCAT- and/or MECR-deficient C2C12, H9c2, and HeLa cell lines. Data are representative of 3 biological replicates. (B) Representative total ion chromatograms of non-treated LA-d_5_ ITSD (top panel) and following chemical derivatization (bottom panel). (C) Schematic of chemical derivatization approach using DMED treatment to visualize LA and LA-d_5_ internal standard by mass spectrometry. (D) Representative overlayed extracted ion chromatograms of derivatized LA from the denoted cell lines.

To directly assess how free cellular LA abundance was affected by perturbation of mtFAS activity, we performed liquid chromatography/mass spectrometry (LC/MS) analysis from cell culture extracts. Free LA was initially difficult to detect (**Fig. 1B**). Therefore, to improve detection and enable absolute quantification we used a chemical derivatization method to add a dimethylamino moiety to the carboxyl group of free LA and spiked in 1 μg/mL of stable-isotope deuterium-labeled LA-d_5_ internal standard^24^ (**Fig. 1C**). Following derivatization, free cellular LA levels were detected, albeit at very low levels, with no appreciable differences among the genotypes (**Fig. 1D**). The derivatized LA-d_5_ internal standard was readily detectable following extraction and derivatization (**Supplemental Fig. 1A-H**), and quantification of the average response ratio of LA/LA-d_5_ sum area also showed no difference between control and mtFAS-deficiency among all cell lines (**Supplemental Fig. 1I**). These data corresponded to absolute quantitation in the range of ~20 pmol – 14 nmol/1000 cells across all samples. Together, these findings show that while mtFAS activity is required for protein lipoylation, free cellular LA is maintained independently of *de novo* biosynthesis. Therefore, free cellular LA must be derived from exogenous or dietary sources rather than mtFAS activity.

### 2.2 LA supplementation raises free cellular LA but does not rescue protein lipoylation

Previously reported work has demonstrated that upon addition to cells, LA is rapidly absorbed, catabolized, and/or excreted^19^. The rapid absorption of LA may reflect passive diffusion across the plasma membrane consistent with its lipid-like structure^25^, as well as transport via the Na+/multivitamin transporter (SMVT)/SLC5A6^26^. We therefore wondered whether treatment with exogenous LA was able to sufficiently raise cellular LA levels due to its rapid metabolism and clearance. Thus we next quantified free cellular LA following exogenous supplementation to levels that have previously been examined for their ability to rescue protein lipoylation in cellular models^27–31^. Cells were treated with supraphysiological concentrations of LA for 24 hours and extracts were derivatized and analyzed by LC/MS (**Fig. 2A-C**). Absolute quantitation revealed a dose-dependent increase in free cellular LA in C2C12 (**Fig. 2A**), H9c2 (**Fig. 2B**), and HeLa (**Fig. 2C**) cells, with concentrations reaching ~ 2 - 30 μmol / 1000 cells. This increase was consistent across all cell lines and independent of mtFAS activity (**Fig. 2A-C**). Despite the massive accumulation of intracellular LA, none of the treatments increased protein lipoylation in control cells, nor did they rescue lipoylation in mtFAS-deficient C2C12, H9c2, or HeLa cells (**Fig. 2D-F**), consistent with previous reports^27–31^. Therefore, we conclude that the failure of mammalian cells to scavenge LA upon supplementation is not a result of insufficient accumulation and/or rapid clearance, but results from the lack of the required enzymatic capability. These data further support a model in which free cellular LA and protein-bound LA are two functionally distinct pools that do not influence one another.

**Figure 2.**
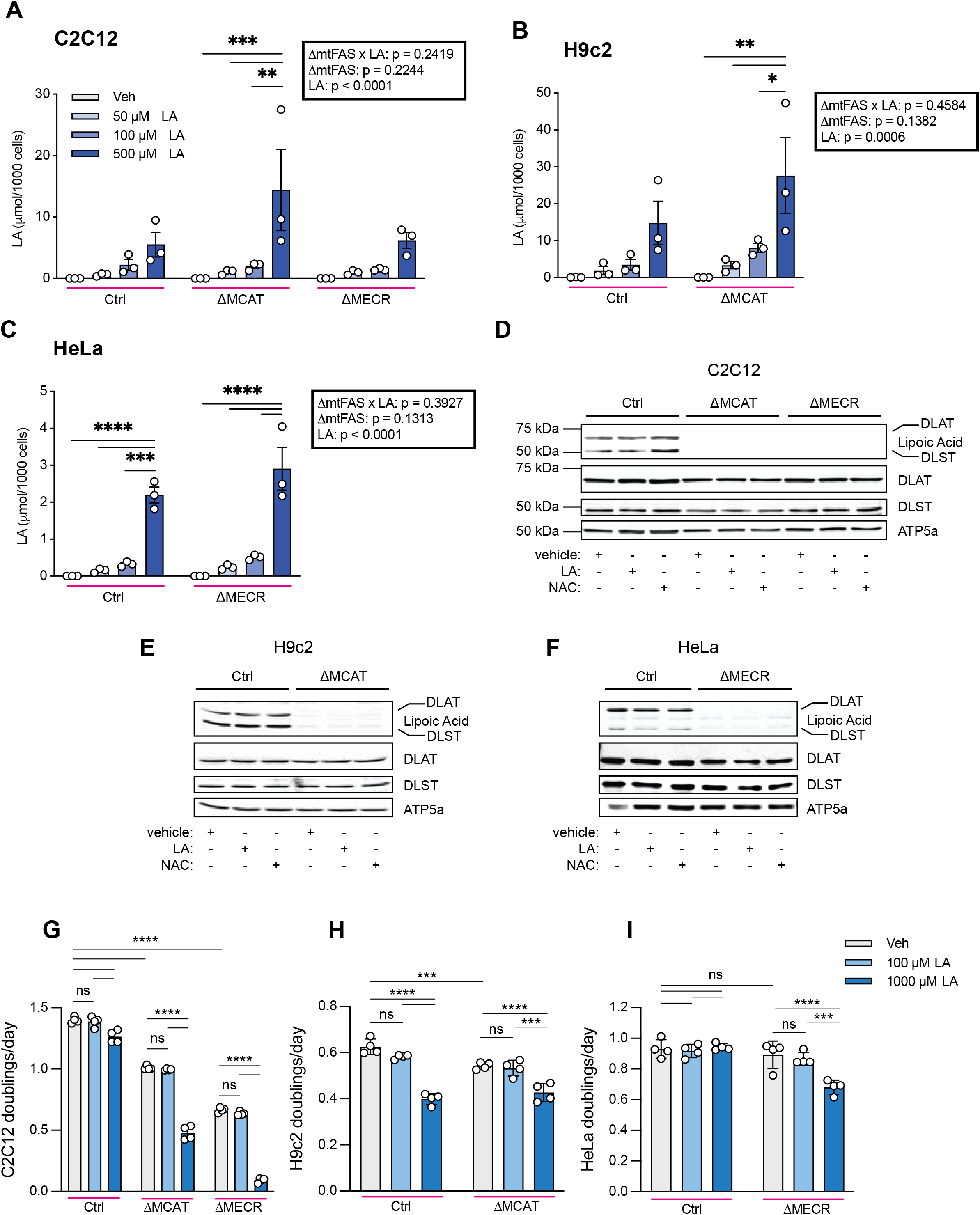
LA supplementation raises free cellular LA but does not rescue protein lipoylation. (A-C) Quantification of intracellular LA from the denoted C2C12 (A), H9c2 (B), and HeLa (C) cell lines supplemented with LA. Data represent the mean LA μmol/1000 cells; error bars denote SEM. * = p <0.05. ** = p < 0.01. *** = p < 0.001. **** = p < 0.0001 as determined by Two-Way ANOVA followed by Tukey’s multiple comparisons test. Statistics for the overall effects of mtFAS deficiency (ΔmtFAS), LA supplementation (LA), and their interaction are denoted in boxed regions. (D-F) Whole cell lysates were separated by SDS-PAGE and immunoblotted for the denoted targets in C2C12 (D), H9c2 (E), and HeLa (F) cell lines supplemented with vehicle, 100 μM LA, or 100 μM NAC. Data are representative of 3 biological replicates. (G-I) C2C12 (G), H9c2 (H), and HeLa (I) cell lines were cultured in an IncuCyte® system with images captured every 4 h for 3-5 days. Data represent the mean doublings per day calculated from technical replicates of growth curves (Supplementary Fig. 2); error bars denote SEM *** = *p* <0.001. **** = p <0.0001 as determined by Two-Way ANOVA followed by Tukey’s multiple comparisons test. Data are representative of 1 experiment from 3 biological replicates.

Given that exogenous LA did not rescue protein lipoylation, we considered whether it may affect cell viability, proliferation, and/or mitochondrial function through alternative mechanisms. LA and its reduced form dihydrolipoic acid (DHLA) have reported antioxidant activity, functioning as metal chelators, free radical scavengers, and regenerators of endogenous antioxidants such as glutathione^19,32^. Additionally, clinical and preclinical studies report beneficial effects of LA on antioxidant activity^17^. We next asked whether LA supplementation could rescue proliferative defects associated with impaired mtFAS activity ^21,33^. LA supplementation did not restore proliferation and instead reduced growth in a dose-dependent manner in control and mtFAS-deficient C2C12 (**Fig. 2G, Sup Fig. 2B-D**), control and MCAT-deficient H9c2 (**Fig. 2H, Sup Fig. 2E and F**), and MECR-deficient HeLa (**Fig. 2I, Sup Fig. 2G and H**) cells. This suggests that mammalian cells exhibit increased sensitivity to elevated LA concentrations under conditions of mitochondrial dysfunction.

### 2.3 OXPHOS function is independent of free cellular LA abundance

In addition to its role in LA biosynthesis, mtFAS regulates iron-sulfur (FeS) cluster biogenesis and ETC complex assembly^22,34,35^. Loss of mtFAS activity therefore results in severe OXPHOS dysfunction that is largely independent of LA biosynthesis, as *Lipt1-*deficient C2C12s largely retain ETC assembly and exhibit comparatively milder OXPHOS dysfunction^22,34–36^. We therefore wondered whether LA supplementation could improve mitochondrial respiration despite having no detectable effects on protein lipoylation, via an alternative mechanism.

Seahorse mitochondrial stress tests were performed on control and mtFAS-deficient cells treated with LA or the antioxidant N-acetyl-L-cysteine (NAC) (**Fig. 3A-F**). In control C2C12 cells, LA supplementation increased basal and ATP-production coupled respiration (**Fig. 3B**). In contrast, LA produced variable reductions in basal, ATP-production coupled, and maximal respiration in H9c2 and HeLa cells (**Fig. 3D and F**). Importantly, LA supplementation did not rescue respiratory capacity in mtFAS-deficient cells across all lines (**Fig. 3A-F**). Similarly, NAC had no effect on protein lipoylation (**Fig. 2E-G**) and increased respiration in C2C12 cells while reducing respiratory parameters in H9c2 and HeLa cells (**Fig. 3B, D, and F**). NAC also failed to restore OXPHOS function in mtFAS-deficient cells (**Fig. 3A-F**). As a whole, these data suggest that OXPHOS function is independent of free cellular LA levels and that any observed benefits from exogenous LA or antioxidant treatment are likely not attributable to improved mitochondrial function.

**Figure 3.**
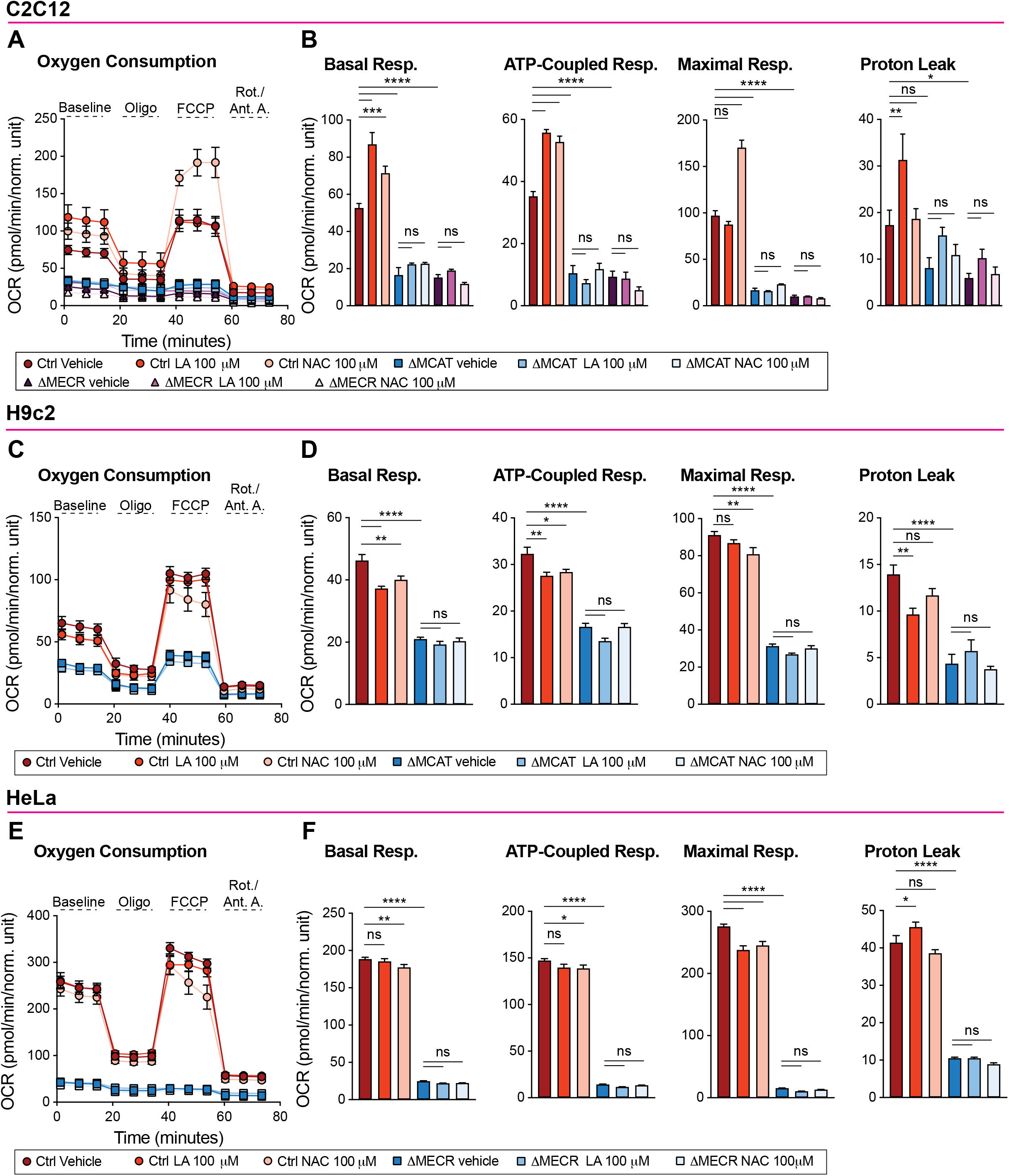
OXPHOS function is independent of free cellular LA abundance. (A-F) Seahorse mitochondrial stress test for Oxygen Consumption Rate (OCR) of C2C12 (A), H9c2 (C), and HeLa (E) cell lines. Data are representative of 1 experiment from 3 biological replicates. Error bars represent SD. (B, D, F) Quantification of basal respiration, ATP-production coupled respiration, maximal respiration, and proton leak from data in (A, C, and E) respectively. Data are mean OCR; error bars denote SEM. * = p <0.05. ** = p < 0.01. *** = p < 0.001. **** = p < 0.0001 as determined by One-Way ANOVA followed by Šidák’s multiple comparisons test.

### 2.4 Elevated free cellular LA buffers ROS insults

LA supplementation has been reported to confer antioxidant benefits in preclinical and clinical studies^17,32^. We therefore directly compared its antioxidant activity to NAC in control and mtFAS-deficient C2C12 cells (**Fig. 4A-F**). Cells were treated with 100 μM LA or NAC and challenged with H_2_O_2_ to induce ROS insult, then H_2_O_2_ levels were assessed using 2’,7’-dichlorodihydrofluorescein diacetate (H_2_DCFDA) fluorescence (**Fig. 4A-C**). Exogenous LA treatment reduced H_2_DCFDA signal relative to H_2_O_2_ alone (**Fig. 4D-F**). However, NAC produced a significantly greater reduction in ROS signal, indicating stronger antioxidant activity compared to LA (**Fig. 4D-F**). These data support the notion that if LA treatment confers any benefits, it is likely via non-specific redox activity. Collectively, our data support a model in which lipoic acid exists in two functionally independent cellular pools: a protein-bound pool generated by mtFAS and required for protein lipoylation, and a free cellular pool derived from exogenous sources with potential antioxidant roles but little effect on mitochondrial function.

**Figure 4.**
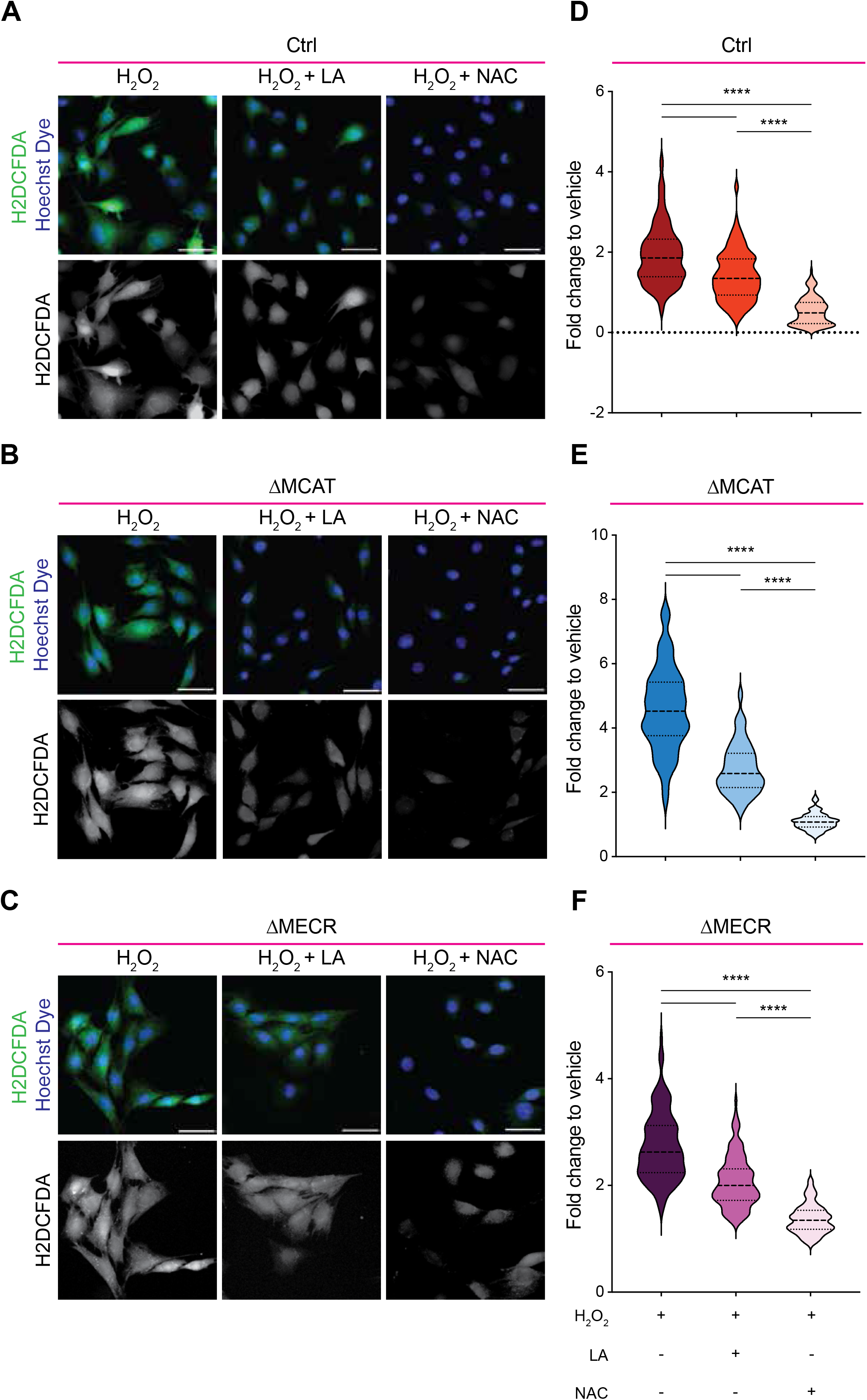
Elevated free cellular LA buffers ROS insults. (A-C) Representative 20X HPF images of control (A), MCAT-deficient (B), and MECR-deficient (C) C2C12 cells supplemented with 100 μM H_2_O_2_ alone and co-treated with either 100 μM α-LA or 100 μM NAC immunostained with Hoechst dye to visualize nuclei and H2DCFDA (green) to visualize ROS. Scale bars = 50 μm. (D-F) Quantification of fold change of mean fluorescent intensity of H2DCFDA expression per cell area (μm^2^) relative to vehicle treated cells in control (H_2_O_2_ *n* = 204, H_2_O_2_ + LA *n* = 147, H_2_O_2_ + NAC *n* = 125) (D), MCAT-deficient (H_2_O_2_ *n* = 145, H_2_O_2_+ LA *n* = 129, H_2_O_2_ + NAC *n* = 139) (E), and MECR-deficient cells (H_2_O_2_ *n* = 127, H_2_O_2_ + LA *n* = 118, H_2_O_2_ + NAC *n* = 116) (F). **** = p < 0.0001 as determined by One-Way ANOVA followed by Dunnett’s multiple comparisons test.

## 3. Discussion

The lack of therapeutic interventions for primary mitochondrial disorders remains a significant clinical challenge. For the past 2-3 decades, “mitochondrial cocktails” have been used to bypass and/or mitigate defects in ETC function; however, their efficacy is difficult to evaluate due to the wide range of compound combinations and clinical heterogeneity, and poor mechanistic understanding of their action^8^. LA is frequently included in these fomulations^17,19,32^. Here, we show that exogenous LA supplementation increases free intracellular LA but does not rescue protein lipoylation or OXPHOS defects in mtFAS-deficient cells, instead exhibiting effects similar to other generic antioxidants. Together, these findings indicate that mitochondrial utilization of LA is independent of cellular LA abundance, raising questions regarding the efficacy of LA supplementation for treatment of primary mitochondrial disorders relative to other cell permeant antioxidant compounds.

No mechanism enabling salvage of environmental LA for protein lipoylation has been identified in mammals, although such pathways exist in prokaryotes^10,15^. Our findings add mechanistic depth to early observations regarding LA supplementation in light of its rapid metabolism, showing for the first time that accumulation of free intracellular LA to μmol levels is possible. Recent studies have demonstrated that mitochondrial expression of the bacterial enzymes Lipoate protein ligase A (*LplA*) or J (*LplJ)* can restore protein lipoylation and improve OXPHOS function of *in vitro* models of LA deficiency upon exogenous LA supplementation^29,30^. Moreover, *LplA* overexpression and LA supplementation was able to rescue embryonic lethality of *Lipt1* KO mice^30^. These findings support a model in which introduction of an LA salvage pathway enables utilization of the free cellular LA pool. In this context, gene therapy approaches designed to confer LA salvage capacity may provide therapeutic benefit in primary mitochondrial disorders characterized by impaired protein lipoylation.

The mtFAS pathway has been cited as the singular source of cellular LA since the discovery that LA was a product of the pathway in 1997 ^37–39^. However, these early studies used extraction methods that would result in liberation of LA from proteins to quantify LA depletion. In fact, LA remains covalently protein-bound during all steps of its synthesis and functional activity as an enzymatic co-factor^15^, and there is no described molecular mechanism to cleave LA from lipoylated proteins to produce free LA downstream of mtFAS. Our data are the first to show that activity of the mtFAS pathway has no bearing on cellular levels of free LA. Together with our observations that exogenous supplementation raises free LA abundance with no effect on protein lipoylation, our analyses suggest a new model in which free LA and mtFAS-derived, protein-bound LA represent two completely distinct LA pools within the cell.

These findings further raise questions about the functional rationale for, and consequence(s) of, maintaining two distinct cellular pools of LA. Exogenous LA supplementation at high concentrations impaired cell proliferation in control cells, with mtFAS-deficient cells exhibiting increased sensitivity to dose (**Fig. 2H-J, Supplemental Fig. 2**). Previous studies have similarly reported proliferative sensitivity to LA supplementation in HEK293T^27^ and in multiple cancer cell models, although the concentrations required to induce these effects was varied^40–45^. Mechanistically, LA-induced cytotoxicity has been associated with increased caspase 3 activity^40^, induction of cyclin-dependent kinase inhibitors^43^, and pro-oxidant activity^44,45^, indicating that free LA can engage in multiple pathways that converge on compromised cell viability. In contrast, recent evidence has demonstrated that lipoylated DLAT, the E2 subunit of pyruvate dehydrogenase, is required for cuproptosis^46^. In this study, Tsvetkov et al. identify protein-bound LA as a key mediator of copper-dependent toxicity and suggest a potential therapeutic vulnerability in tumors enriched for lipoylated proteins. Notably, free LA also binds copper and may serve a protective role against copper toxicity in certain tissues^47^. Together, these observations suggest that compartmentalization of LA into distinct cellular pools may enable differential regulation of metal homeostasis and cellular stress responses. Collectively, these studies and our findings support the need for further work defining the source(s) and role(s) of free cellular LA and its interplay with other cellular pathways and mechanisms involving lipoylated proteins.

## 4. Methods

### 4.1 Cell Culture

C2C12 immortalized mouse skeletal myoblasts (ATCC CRL-1772, verification provided by ATCC), H9c2(2-1) immortalized rat cardiomyocytes (ATCC CRL-1446, verification provided by ATCC), and HeLa (WT and ΔMECR kindly provided by Dr. G Hoxhaj^48^) were cultured in growth media consisting of DMEM with 4.5 g/L glucose, 4 mM glutamine, and sodium pyruvate (Corning, 10-013-CV) with 10% or 20% FBS (Sigma-Aldrich), F0926) at 37°C, 5% CO_2_. For cell culture experiments, cells were supplemented with ethanol vehicle, (±) α-Lipoic Acid (LA, Sigma-Aldrich T1395), or N-acetyl-L-cysteine (NAC, Sigma Aldrich-A9165) at the denoted concentrations and experimental duration. Dialyzed FBS (Sigma-Aldrich F0392) was used for liquid chromatography/mass-spectrometry (LC-MS) and proliferation assays (see below).

#### 4.1.1 Generation of mtFAS mutant cell lines

*Mcat* and *Mecr* knockout C2C12 clonal cell lines were previously generated as described^22^. For generation of mtFAS-deficient H9c2 cells, the sgRNA sequence CTGCCGCTGGATTCAGCGT targeting exon 2 of the rat *Mcat* gene was subcloned into pLentiCRISPRv2^49^. Parental H9c2 were transfected witih pLentiCRISPRv2_rMcat or empty vector using JetOptimus reagent (Polyplus, 76299-630). Approximately 48 hours post-transfection, single GFP+ cells were sorted into 96-well plates to obtain clonal lines. Clonal cell lines were screened by immunoblotting for MCAT protein expression.

### 4.1.2 Proliferation assays

Cells were seeded onto 24-well dishes (Nunc 142475) at densities of 6,000 (Ctrl C2C12), 10,000 (ΔMcat and ΔMecr C2C12; WT HeLa), 12,000 (ΔMECR HeLa), or 5,000 (Ctrl and ΔMcat H9c2) cells per well. After 24 hours, plates were transitioned to an Incucyte® S3 (Sartorius) live cell imager within a humidified incubator at 37°C with 5% CO_2_ for a pre-treatment scan. Plates were then removed, washed once with PBS, then given fresh growth media with the indicated concentrations of LA. The plates were placed back into the Incucyte® and scanned on the phase default setting every 4 hours for 3-5 days depending on when each cell line control reached ~90% confluence. Images were processed using the Sartorius software to define areas of cell coverage and outline individual cells using the ‘Adherent cell by cell’ option. For each cell line, parameters were optimized to ensure accurate cell counting based on distinct morphology. Average cell counts were then exported, normalized, plotted, and the overall proliferation rates were calculated with this formula: doublings per day = (log_2_(final cell count / initial cell count))/days.

### 4.2 Crude mitochondrial isolation

Cells were harvested and processed as previously described^22,33^. Briefly, CP-1 buffer (100 mM KCl, 50 mM Tris-HCl, 2 mM EGTA, pH 7.4) supplemented with mammalian protease inhibitor cocktail (mPIC, Millipore Sigma P8340) was used to resuspend cell pellets. Cells were mechanically lysed and centrifuged at 700 x g for 5 minutes to pellet unlysed cells and debris. Supernatant was transferred to a new microcentrifuge tube and centrifuged at 10,000 x g for 10 minutes to pellet crude mitochondria. Mitochondrial pellets were resuspended in RIPA buffer (10 mM Tris-HCl, 1 mM EDTA, 0.5 mM EGTA, 1% Triton X-100, 0.1% Sodium Deoxycholate, 0.1% SDS, 140 mM NaCl, pH 8.0) with mPIC and used in immunoblotting applications described below.

### 4.3 SDS-PAGE and immunoblotting

Whole cell lysates and crude mitochondrial fractions were normalized for total protein content via Pierce BCA Protein assay (Thermo Fisher Scientific 23225). Protein content was resolved by SDS-PAGE and transferred onto nitrocellulose membranes. Immunoblotting was performed using the denoted primary antibodies (**Table 1**) with visualization of protein signal using Rabbit and Mouse IgG Antibody DyLight 680 or 800 secondary antibodies and a Bio-Rad ChemiDoc Imaging System.

**Table 1.**

| Antibodies | Source and reference number |
| --- | --- |
| ATP5a | Abcam ab14748 |
| Lipoic acid | Millipore 437695 |
| Pyruvate Dehydrogenase E2 (DLAT) | Abcam ab172617 |
| DLST | Cell Signaling 5556 |
| MCAT | Santa Cruz sc-390858 |
| MECR | Proteintech 51027-2-AP |
| Goat anti-Mouse IgG (H+L) Dylight 800 | Rockland Immunochemicals 610-145-002-0.5 |
| Goat anti-Rabbit IgG (H+L) Dylight 800 | Rockland Immunochemicals 611-145-002-0.5 |
| Goat anti-Mouse IgG (H+L) AlexaFluor 680 | Thermo Fisher A-21057 |
| Goat anti-Rabbit IgG (H+L) AlexaFluor 680 | Thermo Fisher A10043 |

### 4.4 Preparation of standard solutions for derivatization of Lipoic Acid

Stock solutions of 2-chloro-1-methylpyridinium iodide (CMPI, 198005-10G, Sigma, 20 µM), N,N-Dimethylethylenediamine (DMED, Sigma 8.03779, 20 µM), and Triethylamine (TEA, Sigma 471283, 20 µM) were prepared in LC/MS grade acetonitrile (Fisher Scientific A955) and stored at −20°C protected from light.

### 4.5 Sample preparation and extraction procedures for Liquid Chromatography – Mass Spectrometry (LC-MS)

Cell samples were prepared by adding 1 mL of normal saline with ascorbic acid (0.1mM) and α-Lipoic Acid-d_5_ (LA-d_5_, MedChemExpress HY-N0492S) (1 ug/mL). Cell plates were scraped and transferred to a fresh tube. Next, 500 µL of ethyl acetate with 10 µL BHT (0.1 mM) and 10 µL formic acid (0.5%) was added to each tube. This solution was vortexed for 30 seconds and centrifuged for 3 minutes. From the top (ethyl acetate) layer containing bioactive lipids, 400 µL was transferred to a fresh autosampler vial (Thermo Fisher Scientific 6PSV9-03FIVP). The extraction procedure was repeated three times for a total of four times. The combined supernatant was dried down under nitrogen at 40°C until samples were dry.

### 4.6 Derivatization procedures

Samples were resuspended in 200 µL LC-MS grade acetonitrile. To each sample, 40 µL TEA (20 µM) and 20 µL CMPI (20 µM) were added and samples were vortexed for 5 minutes. Finally, 40 µL DMED (20 µM) was added and samples were incubated at 40°C for 60 minutes. After incubation, samples were dried under a nitrogen stream. Samples were resuspended in a final volume of 60 µL LC-MS grade acetonitrile (25%). 40 µL of each sample was transferred to a fresh autosampler vial with insert and ran on the instrument.

### 4.7 LC-MS conditions

Samples were analyzed with a Vanquish liquid chromatography system coupled to an Orbitrap ID-X (Thermo Fisher Scientific) using an H-ESI (heated electrospray ionization) source in positive mode. 2 μL of each sample was injected and run through a 9 minute chromatography on a CORTECS T3 column (1.6 μm, 2.1mm × 150mm, Waters 186008500, Eschborn, Germany) combined with a VanGuard pre-column (1.6 μm, 2.1 mm × 5 mm, Waters 186008508). Mobile phase A consisted of 100% LC/MS grade water (Fisher Scientific W6) and 0.1% LC/MS grade formic acid (Fisher Scientific A117-50), and mobile phase B consisted of 99% LC/MS grade acetonitrile (Fisher Scientific A955), 1% LC-MS grade water, 0.1% LC/MS grade formic acid. Column temperature was kept at 40 °C, flow rate was held at 0.3 mL/min, and the chromatography gradient was as follows: 0-1 min from 0% B to 10% B, 1-7.5 min from 10% B to 100% B, 7.5-9 min held at 100% B. A 5 minute wash gradient was run between every injection to flush the column and to re-equilibrate solvent conditions as follows: 0-3 min from 100% B to 0% B at 0.3 mL/min, 3-5 min held at 100% B. Mass spectrometer parameters were: source voltage 3500V, sheath gas 70, aux gas 25, sweep gas 1, ion transfer tube temperature 300°C, and vaporizer temperature 250°C. Targeted acquisition was executed using parallel reaction monitoring (PRM) scans performed in the orbitrap utilizing higher-energy collisional dissociation (HCD) at MS^2^ level to target DMED-derivatized lipoic acid and heavy carbon labeled lipoic acid (see target mass list table below). The quadrupole was used for isolation with an isolation window of 1.6 m/z, orbitrap scan range was 206-284 m/z, RF lens 60%, resolution at 120,000, and HCD collision energy was set to 1. Data dependent MS2 fragmentation was induced in the ion trap with rapid scan rate using normalized HCD collision energy at 30%, the isolation window was 1.6 m/z, and total cycle time was 1 sec.

### 4.8 Targeted metabolomics data analysis

Peak picking and integration were conducted in Skyline (version 26.1.0.057) using accurate mass MS1, MS2 fragmentation pattern matching, and retention time derived from analytical standards run through the described chromatography method. PRM raw data files for all samples of a given experiment were imported and metabolite peaks were auto-integrated based standard verified m/z, precursor adducts, and retention times for all metabolites of interest.

Target Mass list:

| Compound | Formula | Adduct | m/z | Z | t start (min) | t stop (min) |
| --- | --- | --- | --- | --- | --- | --- |
| Lipoic acid (Deriv) | C12H24N2OS2 | +H | 277.1403 | 1 | 2 | 5 |
| Lipoic acid-d<sub>5</sub> (Deriv) | C12H19D5N2OS2 | +H | 282.1717 | 1 | 2 | 6 |

### 4.9 Measurement of mitochondrial oxygen consumption rates

Assays for oxygen consumption were performed on a Seahorse XFe96 Analyzer (Agilent). Briefly, cells were seeded in 96-well plates and cultured overnight with either 0.04% ethanol vehicle, 100 μM LA, or 100 μM NAC. The following day, wells were washed and supplemented with Seahorse DMEM (Agilent 103680) consisting of 25 mM glucose, 2 mM glutamine, and 1 mM sodium pyruvate and either ethanol vehicle, LA, or NAC as denoted.

Standard mitochondrial stress tests were conducted using 1 μM oligomycin, 3 μM FCCP, and 0.5 μM Rotenone + 0.5 μM Antimycin A for C2C12s and H9c2s. For HeLa, mitochondrial stress tests were conducted using 2μM oligomycin, 1 μM FCCP, and 1 μM Rotenone + 1 μM Antimycin A. Measurements were taken over 3 minutes, with three measurements per phase. Data were normalized by measurement of well % confluency using an IncuCyte ® chamber and 10X objective. Results were analyzed in WAVE software and the Agilent Seahorse Analytics browser-based application.

### 4.10 2’,7’-Dichlorofluorescin diacetate (H2DCFDA) staining and live cell imaging

Cells were grown in chambered cover glass slides overnight. Culture media was aspirated, and cells were washed twice in PBS. Serum-free DMEM was then prepared with 2 μM Hoechst Dye and 5μM H2DCFDA and either 0.04% ethanol, 100 μM H_2_O_2_, 100 μM H_2_O_2_ and 100 μM LA, or 100 μM H_2_O_2_ and 100 μM NAC, which was added to respective wells for incubation at room temperature for 30 minutes, protected from light. Media was then aspirated, wells were washed thrice in PBS, then supplemented with HBSS containing 10% FBS and respective ethanol, H_2_O_2_, LA, and NAC treatment conditions. Wells were imaged with a Nikon T*i*2 epifluorescent microscope (20X objective, 0.75 NA).

For quantification of intracellular ROS by 2’,7’-dichlorodihydrofluorescein diacetate (H2DCFDA, Thermo Fisher Scientific D399), regions of interest (ROIs) were manually generated in Fiji software for individual cells, then measured for mean fluorescent intensity (MFI) in the GFP channel. For statistical analysis, outliers from the data set were identified and removed using GraphPad Prism v11 software and the ROUT method, Q = 1%.

### 4.11 Statistics

Statistical analysis was performed using GraphPad Prism v11. Data were analyzed by one- or two-way ANOVA as appropriate followed by two-sided Dunnett’s multiple comparison test (when compared to only control) or Šidák’s or Tukey’s multiple comparison test (when comparing all groups). A p-value of <0.05 was considered statistically significant.

## Acknowledgements

5. Acknowledgments

We thank Dr. Gerta Hoxhaj for kindly providing WT and ΔMECR HeLa cells. This research was funded by the NIH (R35GM151245 to SMN and F30GM154476 to RJW) and the Van Andel Institute (VAI) – Metabolism & Nutrition (MeNu) Program. Stipend support for RJW was provided by the Van Andel Institute Graduate School (VAIGS). We also thank the VAI’s Flow Cytometry Core (RRID:SCR_022685) for their assistance with single cell sorting for generation of clonal H9c2 lines and the Mass Spectrometry Core (RRID:SCR_024903) for their method development for the absolute quantitation of intracellular lipoic acid.

## 6. Author Contributions

Conceptualization (PRN, RJW, SMN), Formal Analysis (PRN, RJW, AEE, MLH), Funding Acquisition (RJW, SMN), Investigation (PRN, RJW, MLH), Methodology (PRN, RJW, AEE, MLH, MRG, RDS), Resources (RDS, SMN), Supervision (RDS, SMN), Validation (PRN, RJW, AEE, MLH, MRG), Visualization (PRN, RJW, MLH), Writing – original draft (PRN, SMN), Writing – review & editing (PRN, RJW, AEE, MLH, MRG, RDS, SMN)

## 8. Data availability

All data needed to evaluate the conclusions in the manuscript are present in the manuscript and/or Supplementary Information. Additional data related to this paper may be requested from the authors.

## 9. Figure Legends

**Supplemental Figure 1.**
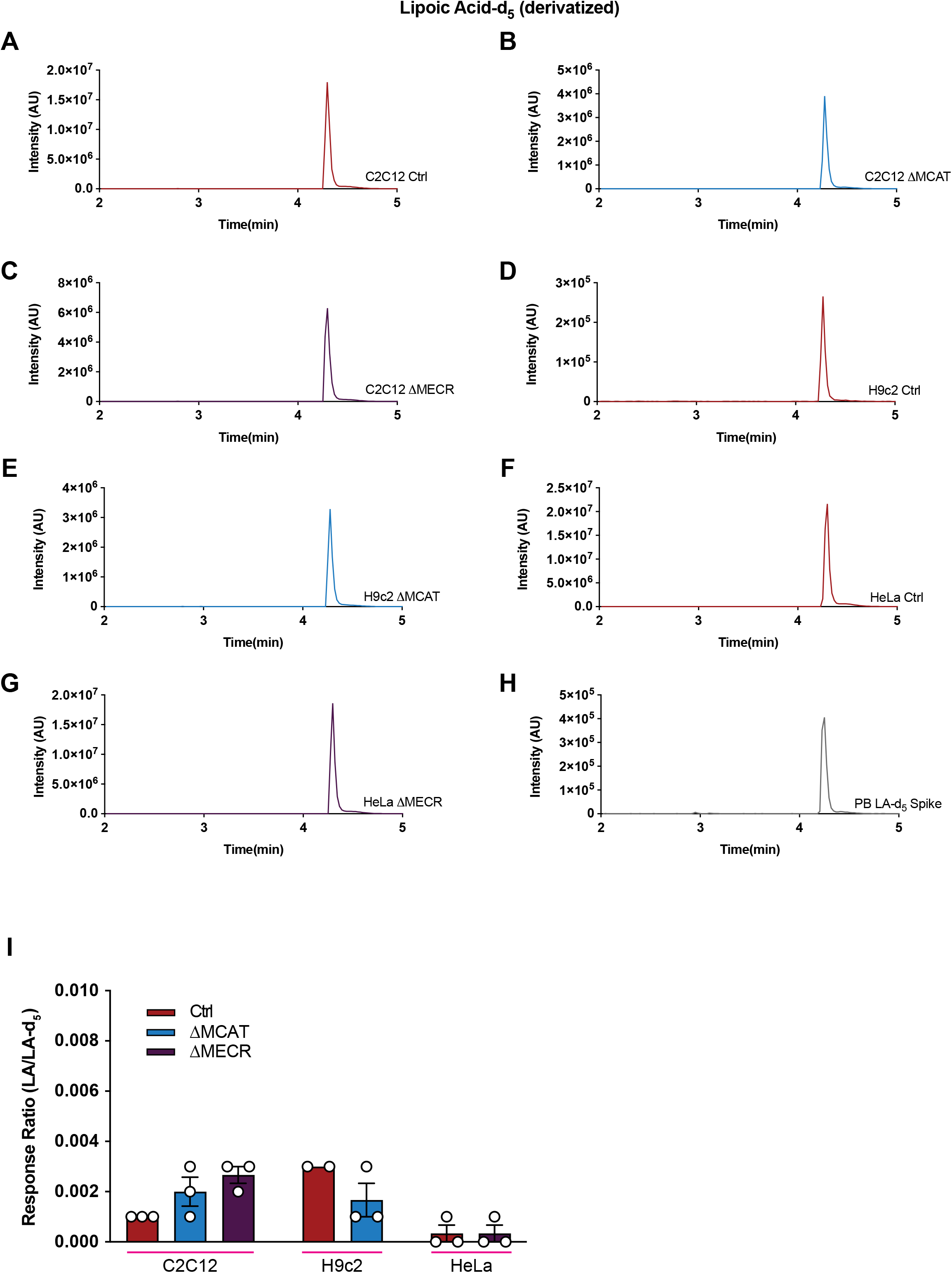
(A-H) Representative extracted ion chromatograms of derivatized LA-d_5_ internal standard from the denoted cell lines. (I) Quantification of the mean response ratio of derivatized LA/LA-d_5_ sum area ± SEM from the denoted cell lines.

**Supplemental Figure 2.**
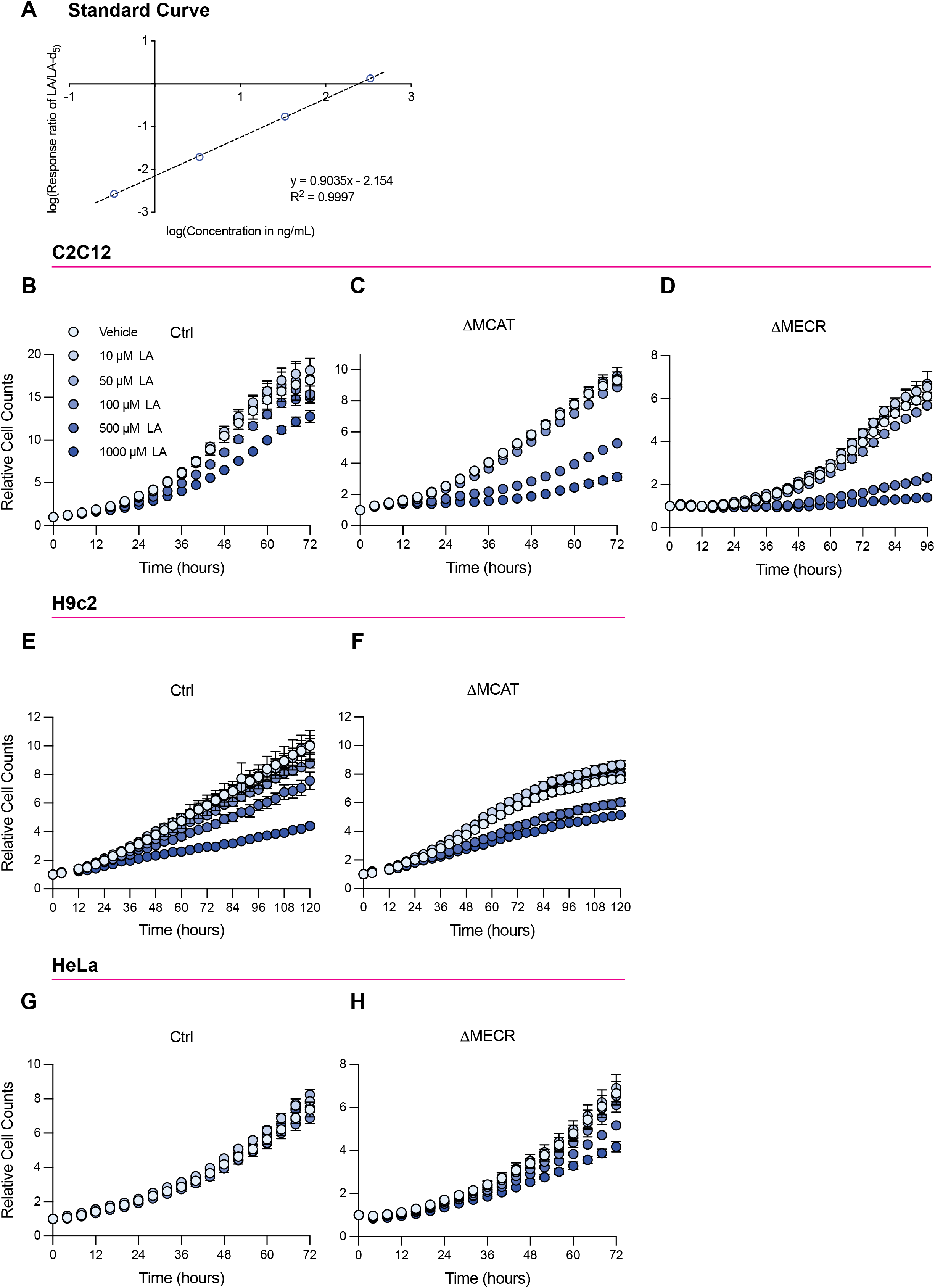
(A) Standard curve of the response ratio of derivatized LA/LA-d_5_ detected by Orbitrap LC-MS. (B-H) Relative counts per well of control (B, E, G), MCAT-deficient (C,F), and MECR-deficient (D,H) C2C12, H9c2, or HeLa cells cultured in an IncuCyte® system supplemented with vehicle or LA for 3-4 days. Data are mean relative cell counts per well; error bars are SEM from n = 4 technical replicates.

